# Genotype-phenotype modeling of light ecotypes in *Prochlorococcus* reveals genomic signatures of ecotypic divergence

**DOI:** 10.1101/2025.10.13.678797

**Authors:** Evan Brenner, Charmie Vang, Connah Johnson, Janani Ravi

**Author notes:** Co-primary authors (Evan Brenner, Charmie Vang).

## Abstract

*Prochlorococcus* species are the most abundant marine photosynthetic bacteria. Despite broadly shared phenotypic traits and marine habitats, they exhibit remarkable genomic diversity. We ask what genomic signatures underlie its ecotypic divergence into high- and low-light adapted lineages, and whether these signatures can still be recovered from incomplete assemblies. From ∼1,000 publicly available *Prochlorococcus* genomes, we focused on those with information on their light adaptation ecotype (high-light/low-light), phylogenetic clades, and depth of isolation. Across these divisions, we calculated average nucleotide identity and constructed pangenomes to assess cyanobacterial core genes vs. those that separate ecotypes. Despite scant conservation, we observe a sharp taxon separation by light ecotypes. Classical machine learning models trained to predict ecotype achieve near-perfect binary classification accuracy even when predicting on partial genomes (Matthews Correlation Coefficient = 0.86 – 1.00), while regression models trained to predict the depth of isolation performed poorly, with high root mean square error values (37.6 – 42.0m). For ecotype prediction, we analyzed top gene features across model runs and classes; these features included photosynthesis-associated genes and pathways, as well as many novel markers of unknown function. When separating ecotypes further by previously described phylogenetic clades, genomic content and composition show even clearer separation among clades, supporting the taxonomic breadth of the *Prochlorococcus* collective. These results emphasize the genomic specialization underlying ecotypic divergence and support the utility of ML approaches for cyanobacterial ecotype prediction from metagenomic data. Expanded sampling will yield novel clade-specific biology. All data, models, and results are available on GitHub: https://github.com/JRaviLab/cyano_adaptation.

**Importance:** *Prochlorococcus* are common aquatic cyanobacteria that can derive energy from light. They can be classified into high-/low-light *ecotypes* depending on how they use light. *Prochlorococcus* have small genomes compared to other bacteria, but the gene sets they carry are also remarkably flexible, which may help them survive and adapt to their harsh oceanic environment. We studied hundreds of *Prochlorococcus* genomes from around the world in an effort to predict ecotypes from partial genome sequences. We used comparative genomics, machine learning, and other statistical methods to identify genomic features associated with ecotypes. These statistical approaches predicted ecotypes accurately, reliably, and according to large differences in gene content and genome structure. Our results support that *Prochlorococcus* can be divided into different species or genera based on clades, and provide many gene targets for further research to understand cyanobacterial circadian rhythms or improve their bioengineering potential as chassis organisms.

## Introduction

*Prochlorococcus*, an ancient cyanobacterial genus^1^, is the most abundant photosynthetic organism in the ocean^2^. Its energy expenditure constitutes a major fraction of global carbon cycling^2^. These bacteria are considered top candidates for supporting the bioeconomy as a chassis organism for bioengineering interventions^3,4^, and are well-known for their ecotypic divergence into high-light (HL) and low-light (LL) adapted clades^2,5–7^. *Prochlorococcus* is also noteworthy for its remarkable genetic repertoire, characterized by a relatively small but flexible genome punctuated by vast numbers and types of accessory genes^8^ — Biller *et al*. reports that every new *Prochlorococcus* strain sequenced adds 160 new genes to a pangenome^4,5^. LL ecotype isolates show more flexibility than HL isolates^5^, but all contain an abundance of rare genes that may only ever be observed in a single isolate and whose functions are generally unknown^4^. Researchers have argued that what was previously called *Prochlorococcus marinus* represents multiple species or even genera that make up a taxonomic collective^5,7–10^. Despite the variability, evidence suggests that the HL/LL division serves as a robust dividing line, suggesting that certain traits may be predictable from genomic content^5^.

This observation underscores a broader challenge in microbial genomics: linking genotypes to phenotypes, especially for uncultivated or environmentally derived taxa^11^. While advances in sequencing yielded an abundance of microbial genomes, linking genomic content to ecological or physiological traits remains complex, particularly when genomes are fragmented, poorly annotated, or lack associated metadata^11,12^. Even with strong phenotypic differentiation, the genomic features underlying traits often remain diffuse and difficult to isolate. This challenge is critical for *Prochlorococcus*: despite being among the best-studied marine bacteria, most sequenced isolates remain poorly annotated and lack associated phenotypic metadata, leaving the genomic basis of ecotypic divergence incompletely resolved.

Genotype-to-phenotype mapping excels at tackling such issues. These methods traditionally rely on comparative genomics approaches such as core and accessory genome analysis, gene enrichment statistics, and targeted gene annotation^11,13,14^. More recently, machine learning (ML) has emerged as a powerful tool for predicting phenotypes directly from genomic data^11,15,16^. These models can incorporate vast numbers of features and capture subtle, nonlinear patterns in genomic variation, offering the potential to link incomplete or draft assemblies with ecological function; this could advance the functional annotation of uncultured, fragmented isolates in metagenomic samples.

Here, we ask *what genomic signatures underlie ecotypic divergence in Prochlorococcus, and can they be recovered from incomplete or metagenomic assemblies?* This question arises because most marine genomic data are fragmentary, and it remains unclear whether such partial genomes still carry enough signal to resolve ecological differentiation. We leverage *Prochlorococcus marinus* as a model organism for microbial niche partitioning^5,6^, applying pangenome-informed ML approaches to connect genotypes to light adaptation phenotypes. We find that classical ML models predict light ecotypes with near-perfect accuracy. This performance holds even when models trained on complete cyanobacterial genomes are applied to fragmented, partial genomes. However, they perform poorly when predicting depth of isolation. These results support that an isolate’s light adaptation ecotype is robustly connected to its conserved genomic content, and that precise environmental isolation source by depth is less detectable through genomic analysis. The wide range of features identified as light predictors support the vast phylogenetic distance others have reported in *Prochlorococcus*. Previously reported phylogenetic clades (described as HL/LL subtypes) were also cleanly separated by their genome composition based on metrics like pangenome makeup, gene coding density, GC-content, or average nucleotide identity. While all genomes used in this research were labeled as *Prochlorococcus marinus* in public databases, their extreme diversity supports that the *Prochlorococcus* collective should be evaluated as separate species or genera, consistent with prior proposals. Novel biology and evolutionary history remain to be discovered in these enigmatic marine bacteria, and substantially expanded environmental metagenome sequencing efforts worldwide can help to resolve these mysteries.

## Results

To assess whether genomic content separates phenotypes like light utilization ecotypes, we analyzed a broad set of publicly available isolates classified as *P. marinus*. However, metadata availability (source: JGI ProPortal) paired with sequenced *Prochlorococcus* genomes (NCBI) constrains most questions in our present dataset (**Figure 1**). We started with the classic and essential adaptation to light (high and low light ecotypes; HL, LL), followed by most likely co-adaptations to depth.

**Figure 1:**
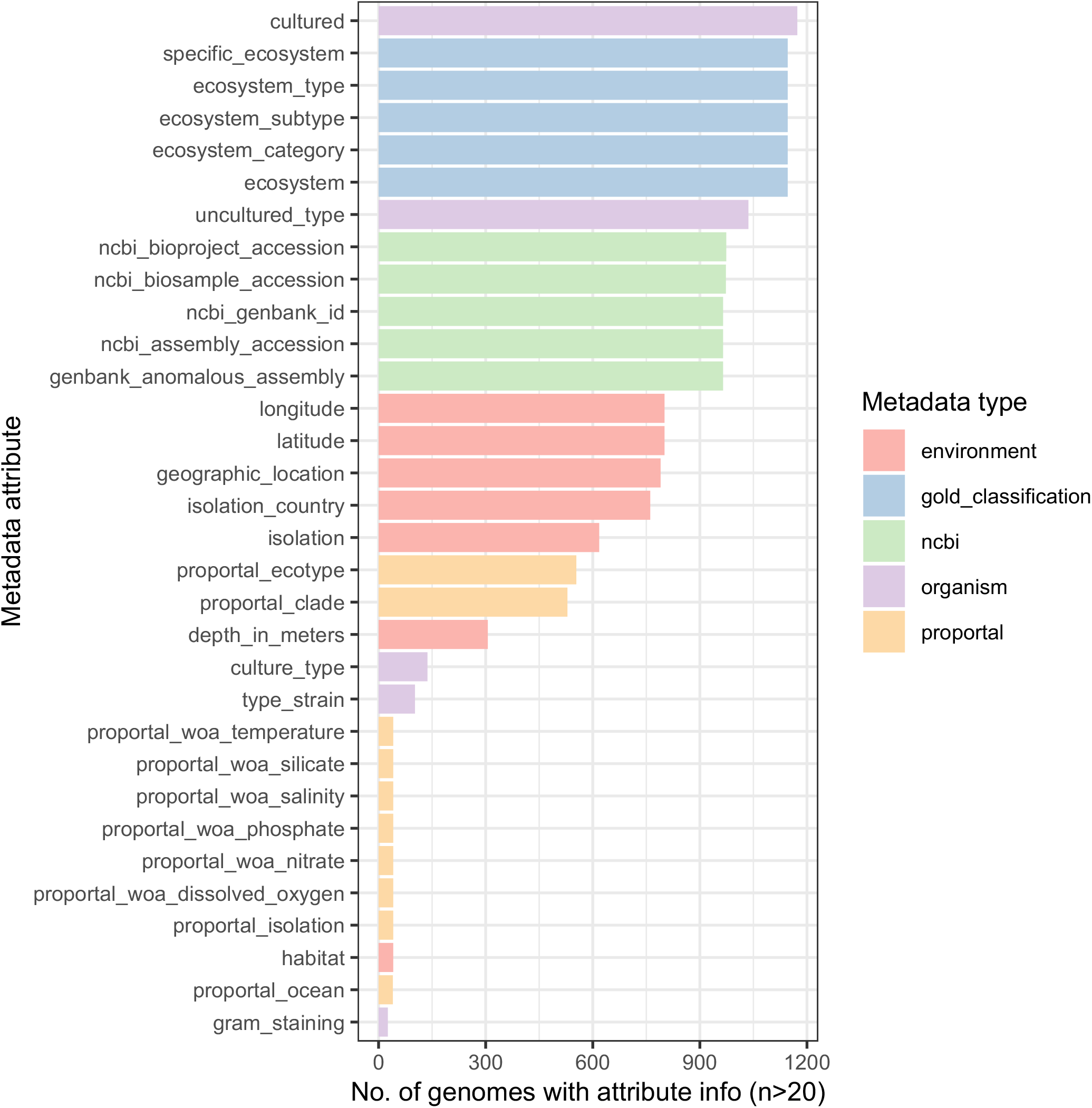
Distribution of metadata availability. In the set of ∼1000 samples with metadata (source: JGI), the proportion of metadata completeness per isolate varied. Metadata field completeness per metadata type (color) and specific attribute (rows) are shown.

We first characterized the pangenome landscape of *Prochlorococcus* to establish the feature space for ML modeling and genotype-to-phenotype mapping. We immediately observed that the primary challenge with *Prochlorococcus* is the lack of a ‘core genome’ by classical pangenome generation tools. Even when we considered only highly complete genomes (BUSCO, see *Methods*) and restricted to HL and LL ecotypes or clades, classical pangenome tools (PanTA, see *Methods*) struggled to capture a core genome due to the extreme sequence and gene content diversity across genomes. Nonetheless, the full pangenome provided a feature set of 78,790 gene groups, with a vast majority (76,764 gene clusters) found in under 15% of genomic isolates.

We sought to capture the overall global structure of genome variations using unsupervised clustering by uniform manifold approximation and projection (UMAP), a dimensionality reduction and visualization technique^17^. When clustering encoded pangenome content per isolate, we found a striking separation of HL and LL ecotypes over UMAP1, even when fragmented genomes were included, suggesting that the ecotype signal is well-embedded in pangenomic variation (**Figure 2**). Given these strongly separated clusters, we observed strong performance across ecotype prediction iterations as expected. ML models consistently successfully predicted LL genomes, but tended to misclassify a minor subset of HL genomes as LL (**Figure 3A**; light blue triangles). These misclassifications became less common as the feature space was reduced for logistic regression (LR) models, with the fewest errors occurring with PCA (principal component analysis) (**Figure 3B**). Interestingly, random forest (RF) models showed zero misclassifications for full and reduced feature spaces, but RF PCA reproduced the misclassification events occurring with other models. The repeated misclassifications restricted to a small, consistent subset of HL genomes may represent transitional or misclassified ecotypes, or may stem from rarer incidents of horizontal gene transfer across ecotypes.

**Figure 2:**
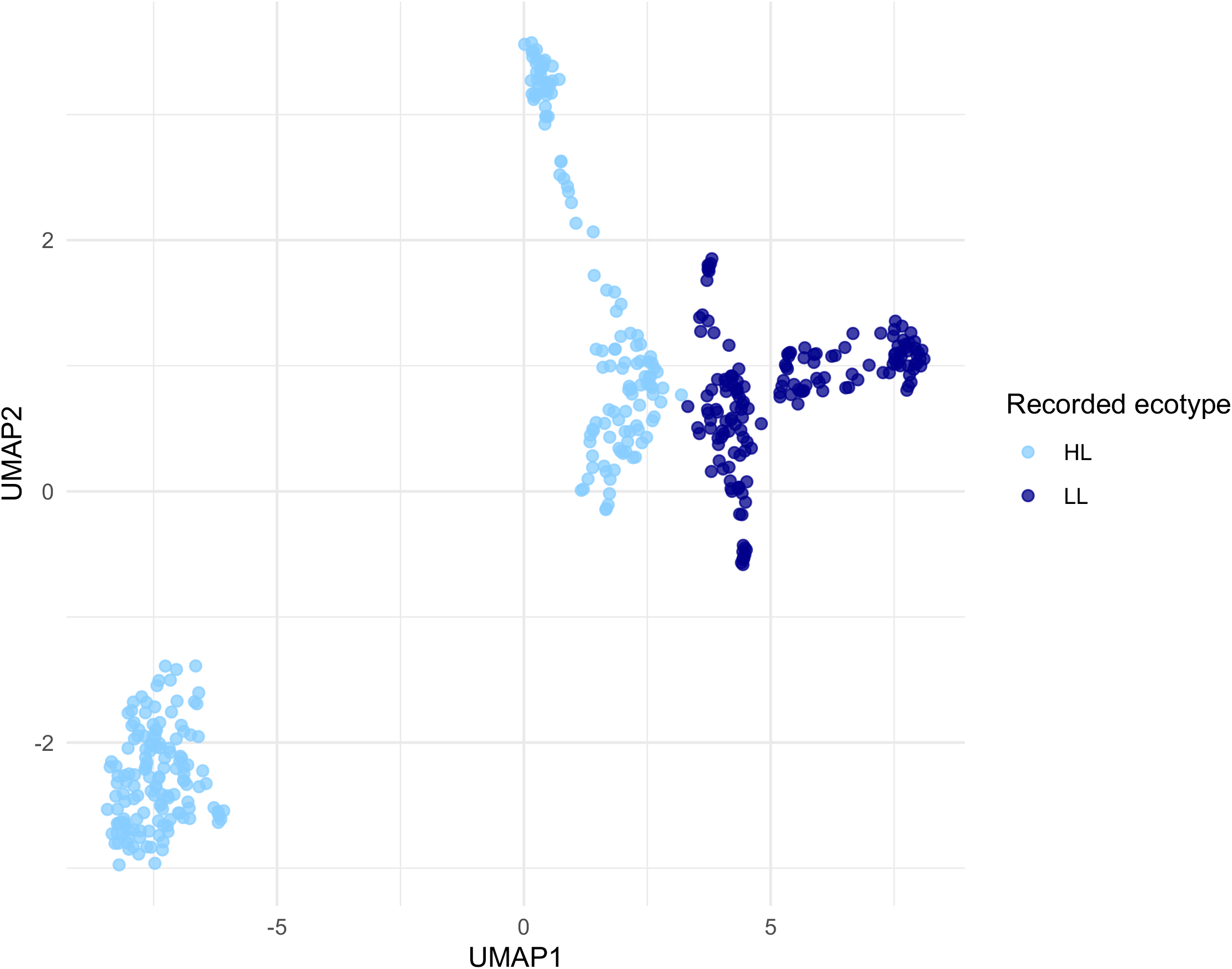
UMAP of genome content for *Prochlorococcus*. We performed unsupervised clustering of the pangenome content of all analyzed isolates and then colored points by their recorded ecotype (high-light, HL, light blue; low-light, LL, dark blue). A strong separation between ecotypes is clear by UMAP1, with distinct, additional subclustering.

**Figure 3:**
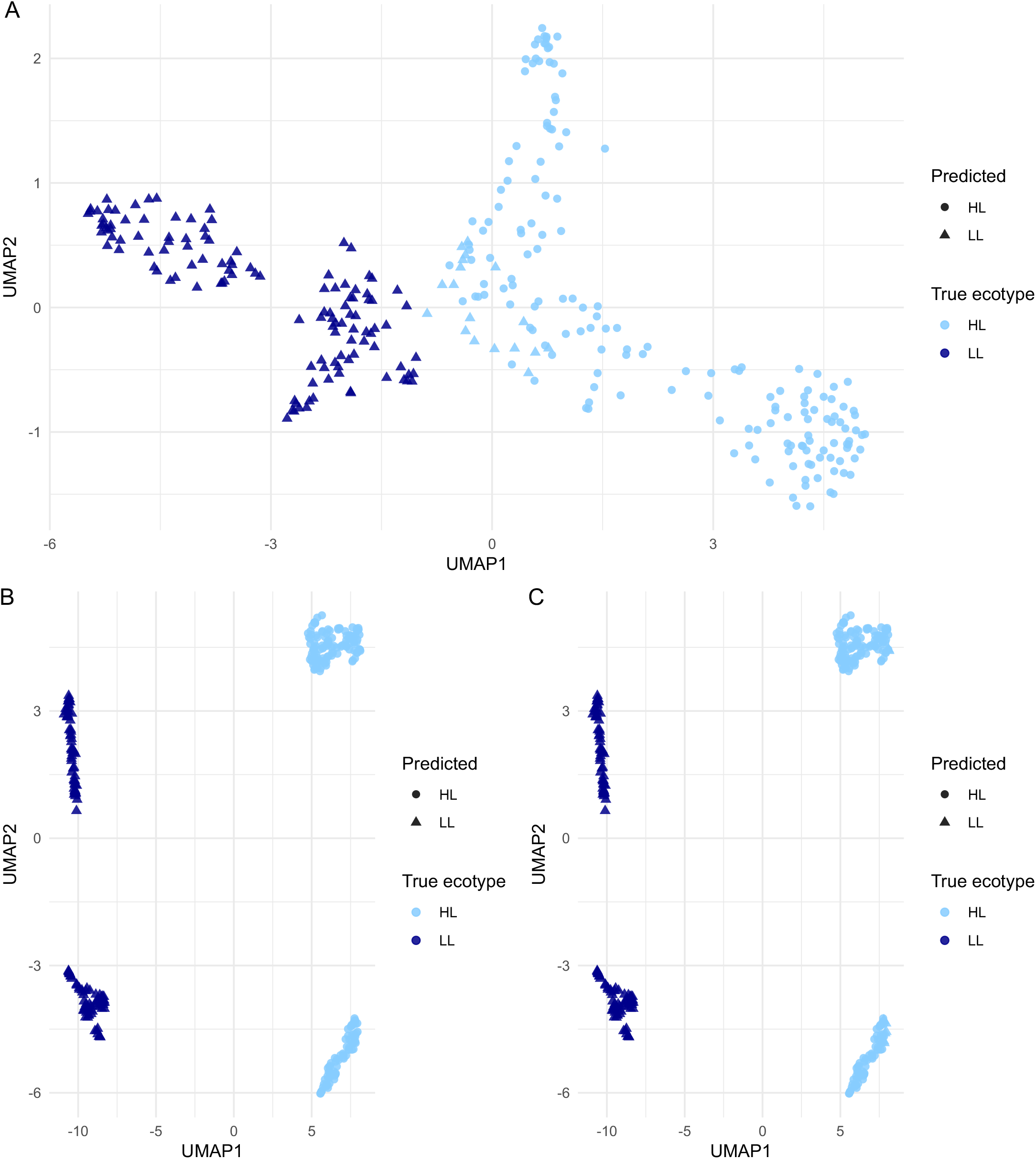
UMAP of test set *Prochlorococcus* genome content with ML classifications. We compared ML model predictions (shapes) with the recorded ecotypes from JGI (colors). **A)** A representative plot from a logistic regression (LR) model run using the full feature space. Misclassifications only occur in one direction — some HL isolates at the intersection between clusters are incorrectly predicted to be LL isolates (light blue triangles, right cluster). **B)** LR model predictions using principal components, PCs, as features, showing no misclassifications and the greatest cluster separation, and **C)** RF model predictions using PCs as features. RF PCA is the only RF model implementation that showed these misclassifications (n=10) despite the strong separation of clusters. Note that genomes in these plots are restricted to those in the test set.

Given the perfect predictive performance of some models, we also tested the robustness of these models using label permutation, in which ecotype labels were randomly shuffled while maintaining the original class balance. These models achieved nearly random performance (LR + RF mean MCC ∼ 0) as expected, but RF yielded striking variance measures across folds (**Figure S1**). To investigate the basis for this high variance, we logged the label permutation results for each iteration and evaluated the percentage of randomly assigned labels that were correct. We found strong-to-extreme correlation (Pearson and Spearman coefficients ≥ 0.9; **Table S1**) between the proportion of labels randomly shuffled to correct categories and permuted model performance. Model performance scaled with the fraction of correctly assigned labels, indicating that even weak residual class structure produces predictive signal. The UMAP separation and performance correlation may reflect both that the two ecotypes are so intensely divergent that even a ∼60/40 shuffle towards correct categories is enough to create modestly predictive models by fitting to extensively ecotype-biased features, even testing against partial genomes. Supporting this, the models with fewest features (PCA, 50 components) showed the lowest performance for these shuffled label models. The separation we see in these UMAPs may reflect both ecological differentiation and underlying phylogenetic structure.

In contrast, depth, a continuous and environmentally variable trait, showed weak genomic signal reflected as poor performance (RF: 37.3m–41.3m; LR: 39.4m–40.3m). Reducing the feature space (Reduced = 6,622 features; PCA = 50 components) marginally improved accuracy. Given that 468 of 490 samples with depth information fell between 0 and 150m, this level of error (∼25% of the typical range) suggests major flaws with any identified predictors. A UMAP colored by depth shows that while HL and LL isolates tend to cluster at higher and lower depths, there is considerable within-cluster variation, with some LL isolates collected near the surface and vice versa (**Figure 4A**). Model predictions of depth, by contrast, are relatively uniform within each cluster (**Figure 4B**).

**Figure 4:**
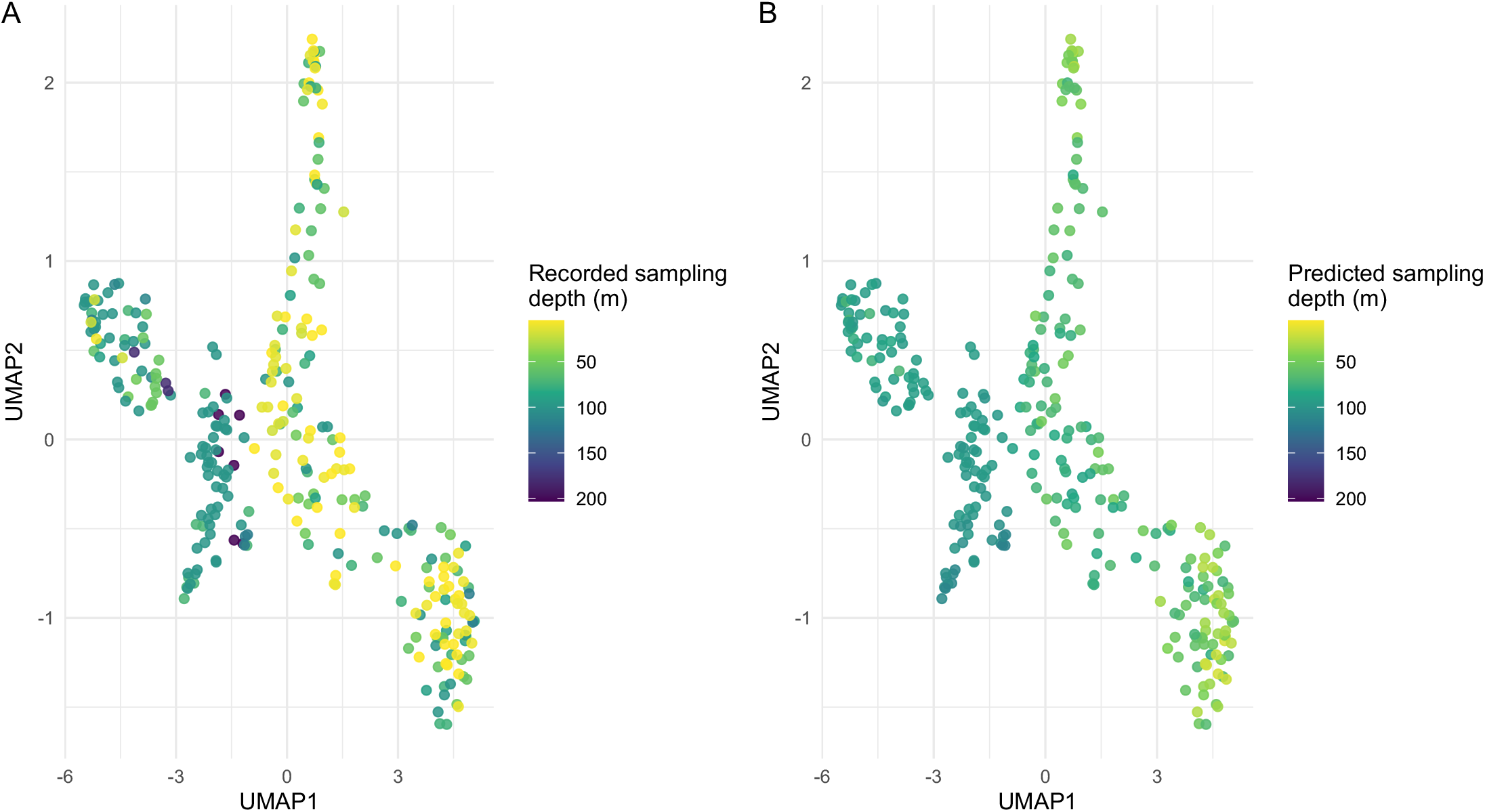
UMAP of test set *Prochlorococcus* genome content colored by depth of isolation. **A)** Clusters were colored by their reported depth of isolation. The left cluster tended to be found at greater depths (corresponding to the LL ecotype cluster), while the right side tended to be isolated closer to the surface (corresponding to the HL ecotype cluster). Variation is observable within and across clusters. **B)** Regression models performed poorly at predicting the depth of isolation, tending to assign fairly uniform values to each cluster. The root mean square error for these models was >20% of the total variation possible in the dataset.

Overall, light-based ecotype prediction models performed exceptionally well. Those using the full and reduced feature spaces performed equivalently when predicting ecotype, but PCA models performed best for LR and worst for RF (**Figure 5**). Across resamples and CV folds, RF tuning using the full pangenome returned perfect performance (MCC = 1.0), suggesting a common fit to highly ecotype-specific genes.

**Figure 5:**
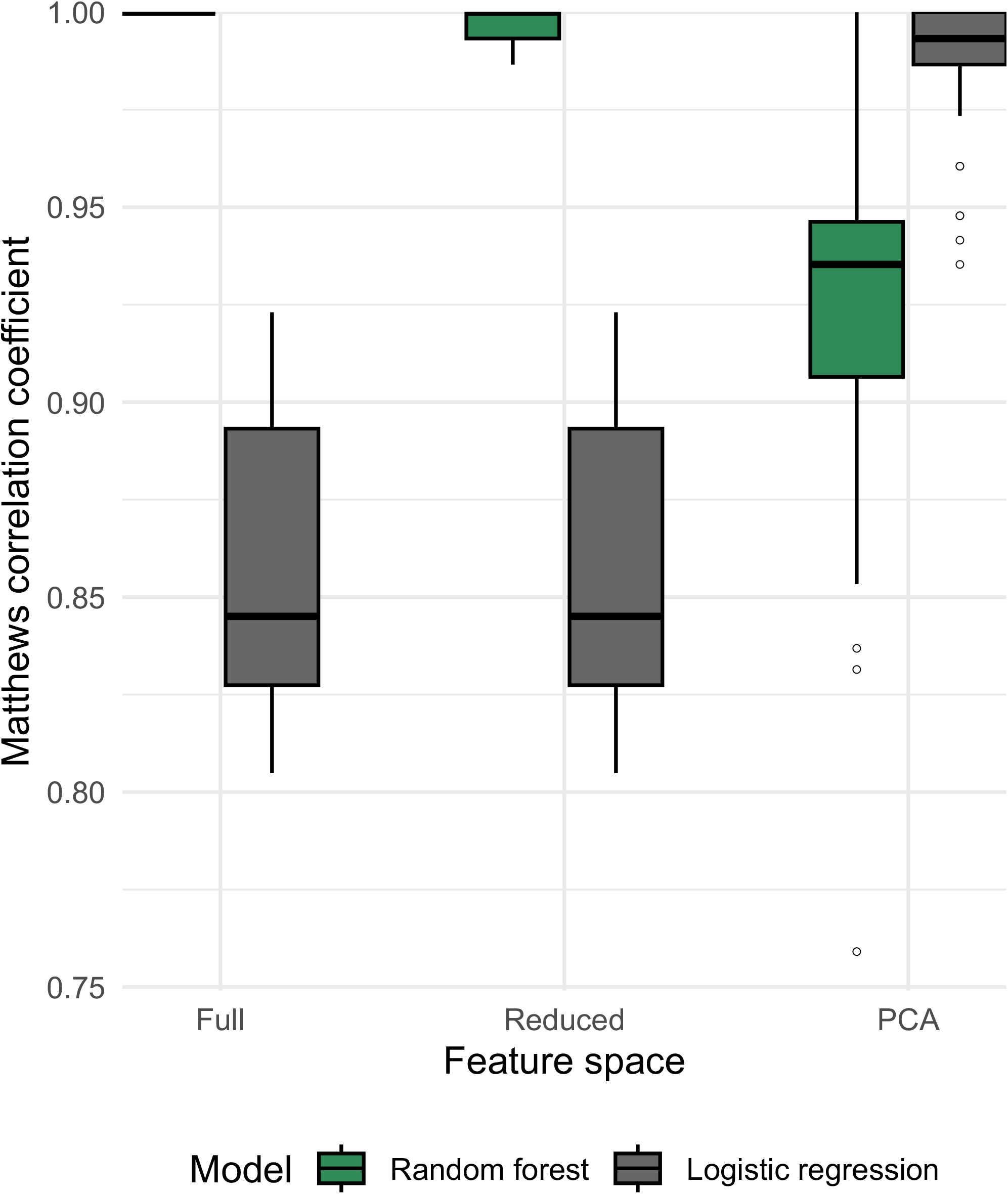
Model performance for binary ecotype prediction. For each model type, 10 randomly seeded models were trained with 5-fold cross-validation, and binary classification performance was recorded for each fold using the Matthews correlation coefficient (MCC). An MCC value of 1 equates to perfect binary classification performance, and 0 is random. We show that while variation exists across the dataset, all algorithms perform exceptionally well. As the feature space is reduced (full → reduced → PCA), LR model performance is improved, while RF performance declines modestly. Shuffled baselines (**Figure S1**) support the performance of these models.

Conversely, we noted very little difference when comparing model types for errors in depth prediction (**Figure 6**). These models showed poor performance regardless of model type (LR, RF) or feature elimination strategy. Shuffled models (**Figure S2**) showed even higher errors, suggesting modeling modestly captures signals of depth adaptation.

**Figure 6:**
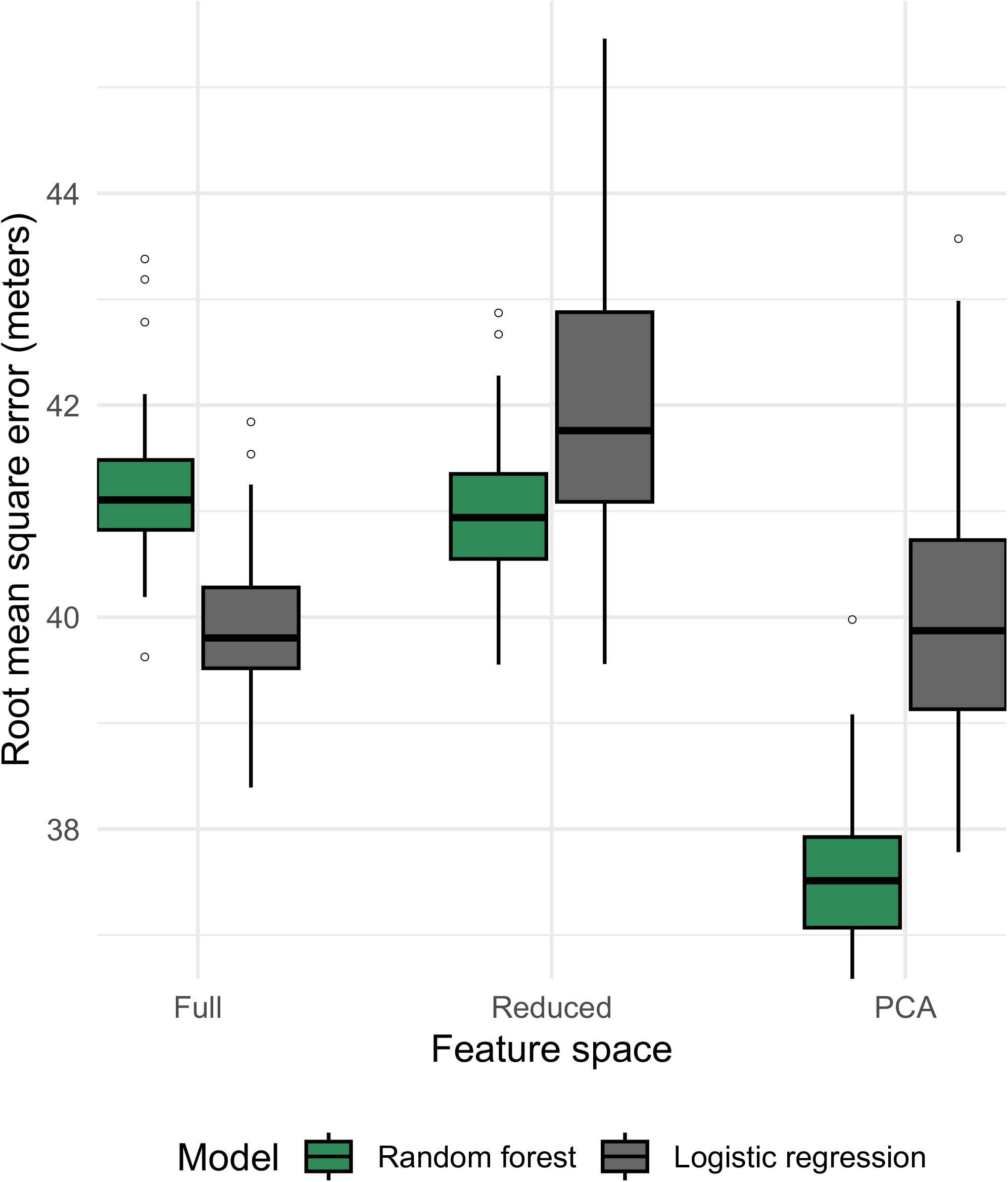
Model performance for regression model depth of isolation prediction. For each model type, 10 randomly seeded models were trained with 5-fold cross-validation, and performance recorded by root mean square error (RMSE) in meters. All models showed high error (∼40m) relative to the overall variation observed in the dataset. No particular model type appeared to do better in this task, suggesting any signal predicting depth of isolation may not be effectively captured by our modeling approach or dataset. Shuffled baselines (**Figure S2**) show these models are marginally improved over random models.

The top predictors driving performance in ecotype classification showed low per-feature importance scores and variation between models, suggesting that performance was not well-explained by any individual genes. Translating high predictive accuracy into biologically meaningful features can challenge ML methods, especially when using large, sparse genomic feature matrices, so we focused our analysis on features that appeared most frequently across 10 randomly seeded model runs.

Many genes were extensively skewed or mutually exclusive between ecotypes. Most top features extracted showed mutually exclusive patterns of ecotype presence/absence. We also frequently observed mutually exclusive feature sets in the reduced feature space models.

Many of these features are consistent with biological understanding of ecotype, including high-light inducible family proteins, UV-B stress response proteins, members and neighbors of the *ndh* operon^18^ and others known or putatively involved in photosynthesis. The gene *pepA* was identified in 94% of HL genomes in training data and 0% of LL genomes, but interestingly, a divergent *pepA* that clustered separately in the pangenome was identified in 95% of LL genomes with 5% prevalence in HL genomes, suggesting a clean evolutionary split between ecotypes that is detectable by differences in this homolog. PepA has been shown to play a role in protection against UV radiation in *Synechococcus*^19^, but any mechanistic differences between the HL and LL PepA remain unclear.

To maximally prioritize robust genetic differences between these ecotypes, we subsetted top features to only those most frequently occurring across model runs and folds, with high feature importance, found in both RF and LR models. This stringent filter yielded just five genes. Among these, *cobI –* which is involved in aerobic corrin ring biosynthesis^20,21^ — was ubiquitous in HL genomes but absent in LL genomes. Intriguingly, *proA*-like genes appeared twice in the top five. Both proteins were exclusive markers for HL genomes in our dataset, suggesting a fundamental role for proline metabolism in the HL lineages. *ndhJ*, an essential member of the NDH-1 complex that facilitates electron flow for the photosystem complex^18,22^, was another shared predictor. Finally, the fifth predictor was a DUF3146 family protein of unknown function widely found across cyanobacteria. Top genes and sequences are shared in **Table S2**.

These striking ecotype differences were also reflected in simple Fisher’s testing (**Figure S3**). Many features (*n*=4,265) were nearly or totally mutually exclusive, returning infinite odds ratios. Of the features shared across ecotypes, 6,283 (7.97% of the total pangenome) were significantly differently distributed between the two ecotypes (adjusted *p* < 0.05; **Table S3**).

We visualized divergence using an all-vs-all FastANI analysis. HL and LL ecotypes formed completely distinct genomic similarity clusters (**Figure 7A**), while depth showed extensive variation within these ecotypes and did not show clear blocks. These findings corroborate the presence of deep ecological and evolutionary separation between HL and LL ecotypes, consistent with prior phylogenetic studies^5–7^. The fragmented nature of many of these genomes means that often only a few regions can be effectively compared, leading to low similarity results. FastANI intentionally does not report ANI values below ∼75% – blank pairs represent a value at or below this point, meaning the true distances between genomes may be greater than what the software is designed to calculate. When analyzing phylogenetic distances by the ITS region, we observed a sharp divide between HL and LL genomes, and while subclades do appear to exist on the tree, they lacked bootstrap support in this analysis (<50) and often showed branch lengths that could not be confidently distinguished from within-clade variation (**Figure S4**). However, clade information has been described previously, and most genomes with ecotype labels also have clade labels (**Figure 1**, “proportal_clade”). We aligned these clade labels to the tree and, despite the lack of bootstrapping significance, saw nearly unanimous agreement of our ITS tree clades with their assigned clades (**Figure S4**). Accordingly, we applied these clade labels to our FastANI comparisons (**Figure 7B**) and UMAP visualizations (**Figure 8**). While some clades like HLII have unexplained separation that may support further taxonomic division, clades generally map well and show good clustering, essentially resolving the FastANI all-vs-all comparisons into neat blocks (**Figure 7B**). The median ANI values within clades (91.1%) vs. between clades (79.8%) were also significantly different by Wilcoxon testing (p << 0.001) Because such clean separation can stem from batch effects and technical artifacts, we checked whether sample BioProject suggested these effects could explain any of the clustering (**Figure S5**). The large volume of data provided by Berube *et al*.^23^ (PRJNA445865) dominated the sample space, but other BioProjects show the same clustering of clades and ANI, supporting that variance is biological and not technical.

**Figure 7:**
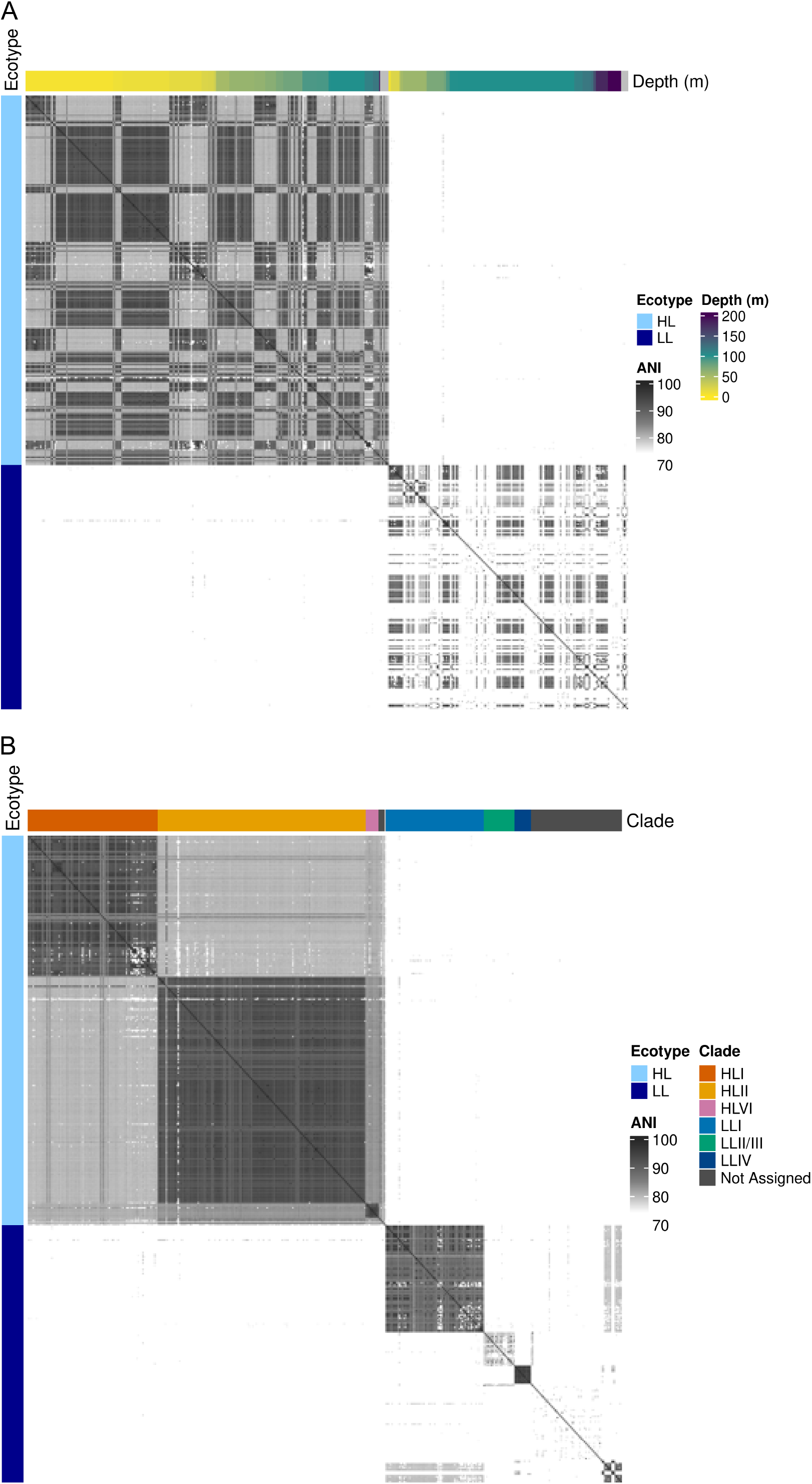
Pairwise average nucleotide identity heatmap of *Prochlorococcus* genomes. We created an all-vs.-all comparison of genomes using average nucleotide identity (using FastANI) and plotted the results as a heatmap. Self-comparisons are along the diagonal (100% ANI, black squares). **A)** The first ordering of comparisons was by ecotype, then depth. Two distinct blocks composed of an HL set of genomes and an LL set of genomes are apparent, though diversity remains within these blocks. **B)** The second ordering of comparisons by ecotype, then assigned clade. This ordering returns more distinct similarity blocks, indicating within-clade genomic ANI is higher.

**Figure 8:**
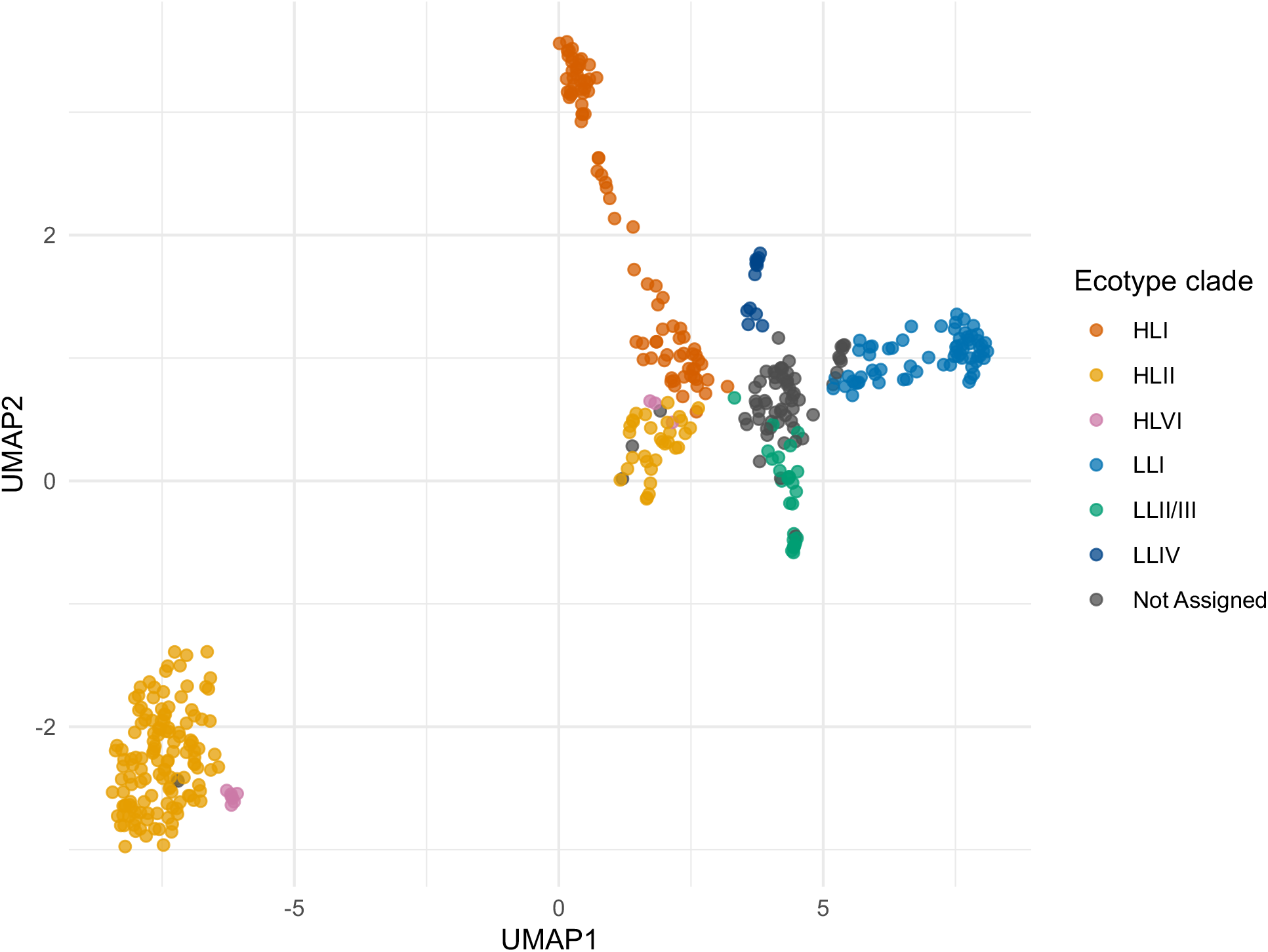
UMAP of genome content aligns well with ecotype clades. When assigned ecotype clade information is overlaid on the UMAP of pangenome content, we see the same HL/LL divide by UMAP1 observed in **Figure 2**, but now also see clades separating for HL and LL clusters largely along UMAP2.

Beyond gene content and ANI, we also observed significant differences between other genome metrics like genome size, %GC content, and coding density across clade categories (**Figure 9**). Statistical comparisons across clades are reported in **Table S4**; however, low *n* for several clades limits our statistical power. For example, clade LLIV shows strikingly different %GC content compared even to other LL clades, but we cannot assign significance with so few high completeness samples.

**Figure 9:**
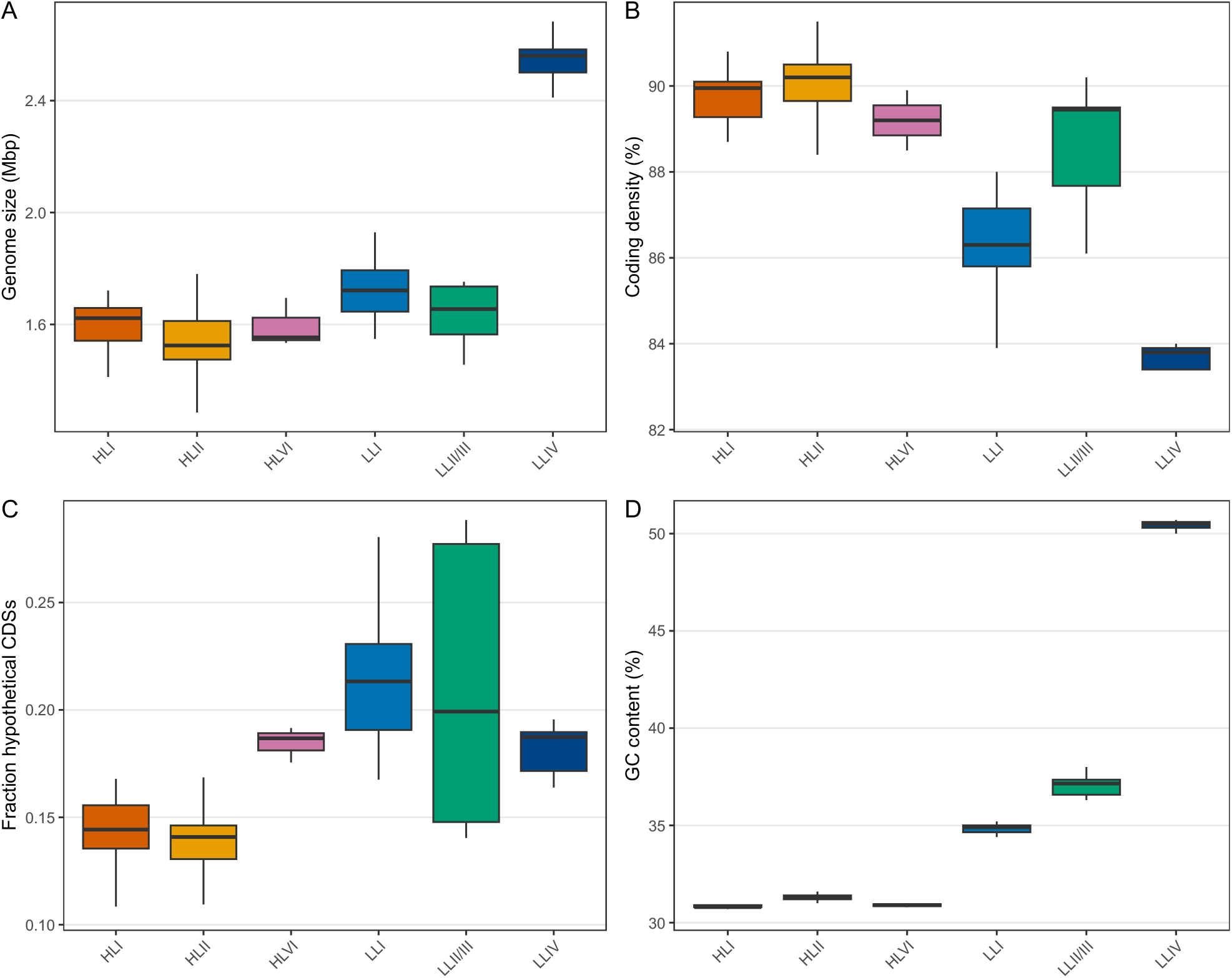
Comparison of genome metrics between clades. We compared basic metrics extracted from genomes (using Bakta) with higher BUSCO completeness (>= 75%) and different clades to discern whether these clades show fundamental differences in genome content. We analyzed (**A**) genome size, (**B**) coding density, (**C**) the fraction of hypothetical genes called, and (**D**) percent GC content. All statistical comparisons are shown in **Table S4**.

In summary, *Prochlorococcus* diversity is unusually extreme among bacteria. Genomics-based classification of ecotype by ML and statistical testing yields excellent accuracy and massively distinct feature spaces, and predictions of ecotype hold well even when models are tested by predicting on incomplete genomes. Predictions for other metadata variables, like depth of isolation, yielded poor performance, suggesting that if genetic predictors of depth as a niche exist, they are not captured by our dataset or analyses. We share our computational workflow through our GitHub repository [https://github.com/jravilab/cyano_adaptation], allowing others to reproduce or extend this analysis.

## Discussion

Our results demonstrate that classical ML models trained on gene presence and absence data can accurately infer ecological phenotypes from draft-quality or metagenomic assemblies. The strong classification performance likely reflects a combination of ecological signal and phylogenetic structure encoded in gene content. The separation in both gene content and nucleotide identity supports that ecotype is a deeply encoded and readily recoverable signal in *Prochlorococcus* genomes^5–7^.

### Prochlorococcus pangenomes are difficult to capture

Pangenomes including multiple ecotypes in this study reliably have very few core genes, even when subsetted to genomes with BUSCO completeness over 75%. We find that current pangenome clustering methods are inadequate to capture shared features across these cyanobacterial isolates, but HMM-based orthology detection^24^ may be better suited for this. While core genomes for *Prochlorococcus* are well-established^25^, and pangenomes built from small (n=12) numbers of isolates in the same clades and with completeness over 80% do return an expected ∼1000 core genes, this signal fades rapidly when incorporating more genomes and more fragmentation. Collectively, this is likely due to both the extensive, known biological variation from one isolate to the next, and the scarcity of near-complete *Prochlorococcus* genomes adding noise to estimates of gene conservation. Additional efforts to improve sequencing depth and coverage will clarify our view of genetic variation across ecotypes, clades, and environmental niches.

### Genetic predictors of ecotype align with biology

Despite the dilution of signal in our gene matrices by the fragmentation of gene homologs into many separate clusters, predictive performance remains excellent. Most of the top gene predictors driving ecotype model performance have apparent roles in differentiating high- and low-light lifestyles: a putative high-light inducible protein with a light-harvesting complex domain (PFAM00504); a hypothetical protein Ycf35 suspected to play a role in photosynthesis (containing DUF1257); the photosynthesis-essential *obgE*^26^, induced by UV-B stress and protection; and even an M3 family peptidase encoded directly adjacent to the photosystem-supporting Ndh-1 complex member *ndhG*^18^. When subsetting further to only top features returned by both LR and RF models, we continue to see generally expected results in the genes returned. In this pool and in each of the larger LR or RF sets, we also regularly see hypothetical proteins with no known or predicted function. A DUF3146 family protein (cluster ID ‘*groups_468*’ in the pangenome; **Table S2**) is ubiquitous across Cyanobacteriota, yet here serves as a strong dividing line for HL and LL ecotypes within *Prochlorococcus*. In the wider set of LR features, a DUF2997 domain-containing protein (‘*groups_586’;* **Table S2**) is most common in cyanobacteria, but also found across other bacterial phyla and even into archaea, eukaryote, and some viruses. Hypothetical and sporadically conserved genes like these unnamed groups/DUF domains are a major fraction of cyanobacterial diversity, and while our results suggest contributions to differentiating clades, ecotypes, and light adaptation, functional studies are necessary to determine their roles. It is also important to recognize that the highly distinct clades may be dominant drivers of separation rather than light-utilization specifically. These *Prochlorococcus* clades and ecotypes offer a trove of uncharacterized biological function in their hypothetical gene space, much of which is unique to cyanobacteria and the hostile oceanic niches they dominate.

### Genomic variability within Prochlorococcus supports taxonomic expansion

Most striking among these results is the extent of separation between high- and low-light ecotypes. Our results across methods, including UMAP, supervised machine learning, average nucleotide identity, genome metrics, and even simple Fisher’s testing, show such a sharp divide between ecotypes and across clades that it seems probable that *Prochlorococcus marinus* is multiple species or genera. Indeed, prior work found similar distances and supported separation, including Thompson *et al*. first in 2013^9^ and more recently from Tschoeke *et al*. in 2020^10^. While global *Prochlorococcus* sampling efforts have been underway for decades (**Figure S6**), we must recognize that ∼500 isolates can capture only a fraction of biological variability. For instance, the single sampling cluster in the Indian Ocean in **Fig. S6** reminds us of the relative sparsity of available data. We expect that expanded sequencing efforts and deeper study of the members of the *Prochlorococcus* collective will reveal new frontiers of biological discovery. These vastly divergent organisms encode evolutionary trajectories for the development of circadian regulation in cyanobacteria, unparalleled genetic variation across tens of thousands of unstudied genes even by the small sample set studied here, and the potential for powerful metabolic pathway rewiring towards enhanced bioproduction^27–29^.

### Clades of the Prochlorococcus collective are separable genotypically and phenotypically

Together, our results indicate that ecotype is encoded as a highly redundant, lineage-associated genomic signal distributed across many gene features. Given the lack of conserved core genes identified in our pangenome, a refined assignment of homology for the extensive gene space would facilitate better interrogation of ecotype- or clade-specific genes. Our ML models’ predictive performance undoubtedly reflects more granular *Prochlorococcus* clade structures rather than solely HL or LL ecotypes. However, separation by these genes does support the previously observed evolutionary divergence and specialization of *Prochlorococcus* ecotypes and of the collective^5,7,9,10^. While the minimal gene sets discussed here are far from exhaustive, we show that the vast library of cyanobacterial genes without known function could be associated with specific phenotypes and traits using ML and statistical approaches. Towards this goal of characterizing the unknome^30^, supporting and expanding ongoing sampling efforts for cyanobacteria, developing new methodologies to better cluster and differentiate the complex pangenome space of *Prochlorococcus*, and investigating novel approaches to expand on the evolutionary trajectory of this phylogenetically complex taxonomic collective are necessary directions to advance research into cyanobacteria and beyond.

## Methods

### Overall data curation, processing, and analysis

We designed our workflow in R and RStudio, with external software dependencies invoked through Docker^31^/Singularity^32^ container calls. We used containers from the open-source State Public Health Bioinformatics Group’s extensive Docker Hub collection^33^. In brief, we acquired genome data and metadata from the Department of Energy’s now-deprecated ProPortal database built from their IMG infrastructure^34,35^, and NCBI through their Datasets CLI^36,37^. We analyzed genomes for completeness through BUSCO^24^, annotated with Bakta^38^, built a pangenome using PanTA^39^, compared genome-genome similarity with FastANI^40^ and RAxML-NG^41^, and performed most other analyses in R. Our work relied on numerous R packages: ggplot2^42^, ComplexHeatmap^43^, magick^44^, circlize^45^, dplyr^46^, readr^47^, stringr^48^, tidyr^49^, tibble^50^, viridisLite^51^, here^52^, tidyverse^53^, jsonlite^54^, purrr^55^, tidymodels^56^, vip^57^, doFuture^58^, future^59^, rstatix^60^, and uwot^61^. Heavy data processing tasks were performed on the University of Colorado’s Alpine HPC cluster using SLURM^62^. For feature investigation, we used NCBI’s BLAST^63^ and our MolEvolvR web-app^64^. Our workflow is available via our GitHub repository [https://github.com/jravilab/cyano_adaptation] and provides a reproducible, end-to-end workflow from genomic acquisition and pangenome construction, ML modeling, and visualization, enabling application to other microbial system. Detailed methods are provided below and in our repository.

### Curating cyanobacterial genomes and phenotypes

We curated cyanobacterial genomes from the DOE-funded Joint Genome Institute^65^ (JGI), of which 503 had light ecotype metadata (categorized as high-light or low-light), 490 of which also included depth of isolation. We retrieved raw genomic assemblies via the NCBI Datasets command-line utility^36^ using the provided NCBI assembly accession IDs. Many assemblies failed to pass through NCBI’s PGAP, prokaryotic gene annotation pipeline^66^, or contained minimal annotation, prompting us to re-annotate all genomes using Bakta (v11.11.4, database v6.0, parameters = --compliant --genus Prochlorococcus --species marinus) to ensure standardized gene prediction^38^.

### Comparative genomics

To evaluate genome quality, we ran BUSCO (v6.0.0) against the cyanobacteria lineage dataset (cyanobacteriota_odb12) and recorded their per-genome BUSCO values, referred to herein as completeness scores^24^. Genomes with >25% completeness (n = 437) were used to construct a pangenome using PanTA, a rapid gene count matrix generation tool^39^. A full pangenome matrix was constructed across all training and testing genomes. Assemblies with higher completeness (>75%; n = 129) were labeled for ML model training, while those with intermediate scores (25–75%; n = 308) were used for testing—genomes with <25% completeness were not used for this analysis.

### Machine learning model setup

To predict light phenotypes, we trained random forest (RF) and logistic regression (LR) classifiers on the *Prochlorococcus* pangenome matrix (features were binary presence/absence of genes) using ranger^67^ and glmnet^68–70^. To assess the robustness of our approach, we trained the models only on high-completeness (>75% completeness) genomes and then applied them to the lower-quality test set (25%–75% completeness). Models were trained using 5-fold cross-validation (CV), and 10 randomly seeded resamples. Compared to their original ∼60:40 HL:LL class balance distributions, a downsampling approach for 50:50 HL:LL class distribution yielded no changes to model performance. Distributions in train and test folds were therefore maintained at their original ∼60:40 HL:LL splits for simplicity.

### Feature space reduction

Given the extreme feature space size (78,790 genes) in contrast to ∼500 isolates, and an overwhelming abundance of genes identified in only one or two genomes (expected in *Prochlorococcus*^5^), we constructed two additional reduced feature matrices by creating a list of genes present in <5% of both HL and LL subsets – if the frequency was below this cutoff in both ecotypes, the gene was removed from the reduced pangenome. Subsequently, the tidymodels^56^ recipe implementation of principal component analysis, PCA (num_comp=50) was used on the reduced feature space to further reduce the number of features. These two reduced subsets were used for modeling separately through the same workflow as the whole feature space to test whether dimensionality reduction was necessary for reliable performance or feature interpretation.

### Binary ecotype classification modeling

Ecotype models were optimized for Matthews correlation coefficient (a robust metric for binary classification with imbalanced classes^71^) and then applied to test data. For each fold in each resample (n=50), we also independently saved and tested the fit on the test data and recorded metrics to assess performance and variance across permutations. We recorded the frequencies at which specific features appeared as top predictors across permutations and their importance scores, and focused only on those features that showed consistent, high signal. Mean, median, and standard deviation of MCC across all permutations were recorded.

### Depth of isolation regression modeling

For regression models that predict sample origin depth in meters, we report performance root mean square error (RMSE) to quantify the difference between predicted and actual values.

### Label shuffled permutation models

Additionally, to test robustness, we also implemented random label shuffling. For each ecotype permutation, per-isolate HL/LL labels were reordered at random while maintaining the overall class balance ratios. For depth models, the depth of isolation values were similarly randomly shuffled. These shuffled label models were run with 5-fold CV, and repeated 10 times. All models were evaluated to provide baseline performance values when the connection between genotype and phenotype was severed.

### Basic statistical analysis

We sought to determine simple statistical differences between features in HL and LL pangenomes. We performed Fisher’s exact tests and adjusted for multiple comparisons. After separating by ecotype clades (e.g., HLII, LLIV), we collected genome metrics from Bakta output files, and performed Wilcoxon rank sum testing for ecotype and Kruskal-Wallis testing for clades to assess whether significant differences existed. We performed post-hoc Dunn testing with Benjamini-Hochberg correction to define differences between specific categories.

### Unsupervised learning and sequence similarity-based clustering

Using the uwot^61^ package in R^72^, we clustered per-isolate pangenome content using UMAP dimensionality reduction and overlaid ecotypes, clades, depth, and BUSCO score. In parallel, we conducted an all-vs-all genomic similarity analysis using FastANI (v1.34) to examine overall nucleotide-level divergence across ecotypes and assess whether HL and LL ecotypes and clades separate consistently in alignment-free space^40^.

### Interpreting key features

We assessed feature importance values for each model run using the vip R package^57^. Features that repeatedly appeared within multiple model iterations or across multiple model types were considered higher confidence. We processed representative protein sequences for genes of interest into domains and motifs using MolEvolvR^64^, and queried for homologs using BLAST^63^.

### Phylogenetic clustering

To compare the diversity of isolates by traditional metrics, we extracted the internal transcribed spacer region between 16S and 23S rRNA genes from our Bakta GFF and FNA files. Given the variable completeness of the genomes in this study, we only extracted sequence if the 16S and 23S rRNA genes were intact and on the same contig, which yielded 339 ITS sequences. We aligned these sequences using MAFFT^73^ (v7.526; options: --localpair --maxiterate 1000), ran ModelTest-NG^74^ (v0.1.7), and determined that the TPM3uf+I+G4 model was best for modeling. We ran RAxML-NG^41^ (v1.2.1) using this model and 1,000 bootstraps to generate a maximum likelihood phylogenetic tree.

## Acknowledgments

This work was enabled through collaboration and discussion with the NW-BRaVE teams at Pacific Northwest National Laboratory (PNNL); we would like to particularly thank Drs. Ruonan Wu, Margaret Cheung, and David Pollock.

