## Supplementary material for "Genotype-phenotype modeling of light ecotypes in *Prochlorococcus* reveals genomic signatures of ecotypic divergence"

**Supplementary Figures** 2

**Supplementary Tables** 8

Supplemental figures

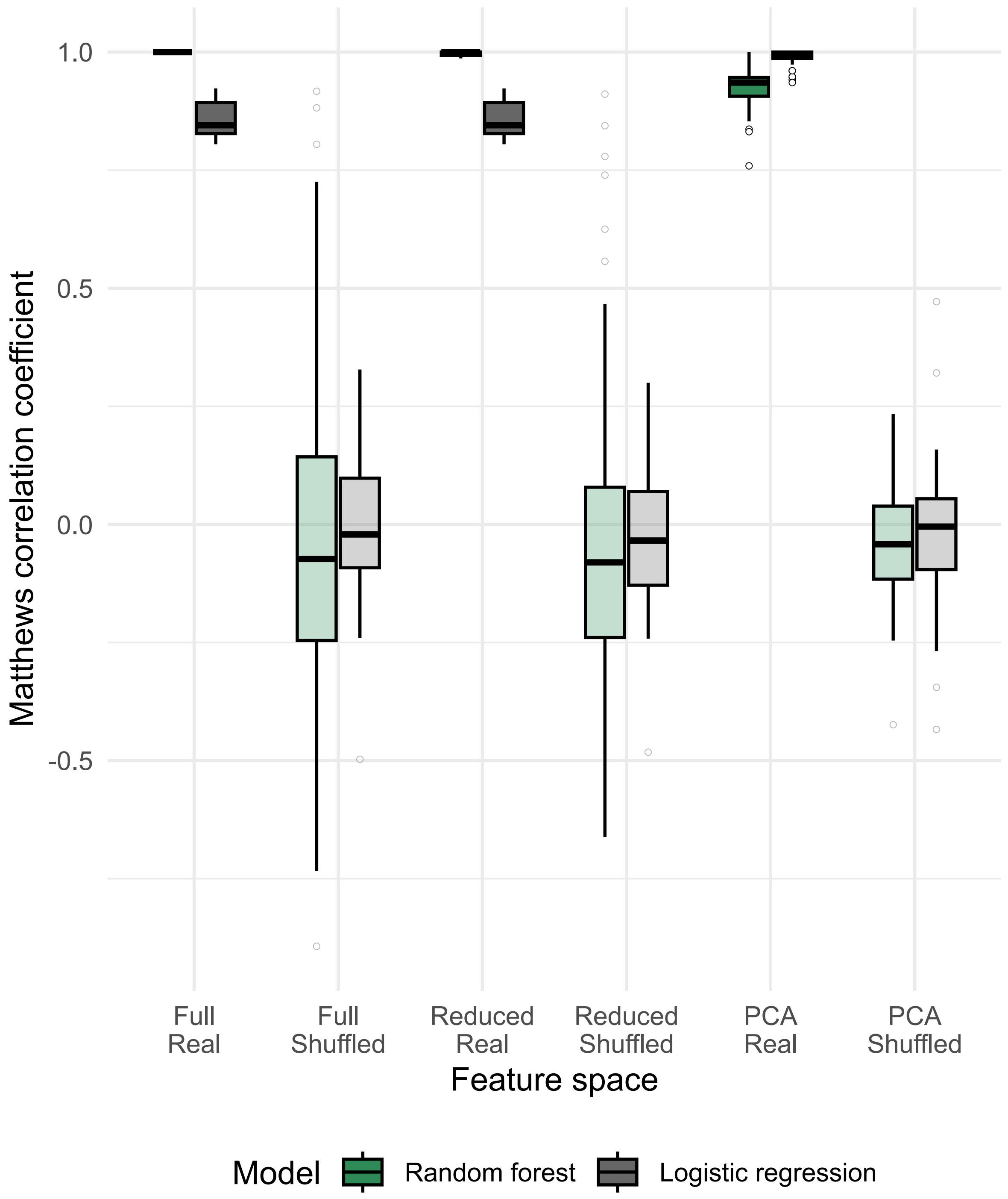


**Figure S1. Ecotype performance vs. shuffled label baselines.**

For the two algorithms chosen (logistic regression and random forest) and three feature spaces used (full pangenome, reduced pangenome, PCA components), we compared classification performance of all folds and resamples runs for the models trained on real data (ecotype labels taken from JGI metadata) and data where the ecotype labels were randomly shuffled. For real models, MCC ranges from 1.0 to 0.86. For shuffled models, median MCC is near 0, indicating expected random performance. However, the MCC range for RF permutations was extremely high for the full feature space and fell to match LR's range as the feature space was reduced. We identified that when randomly shuffled labels approached 60% correct class assignment by chance, these folds became extremely accurate and found a high correlation between random correct class proportionality and MCC in the shuffled models (**Table S1**). The top performance for real RF models, and the propensity for shuffled RF models to pick up minor signals that are accidentally introduced through random shuffling supports that ecotype-specific features cleanly delineate HL/LL ecotypes.

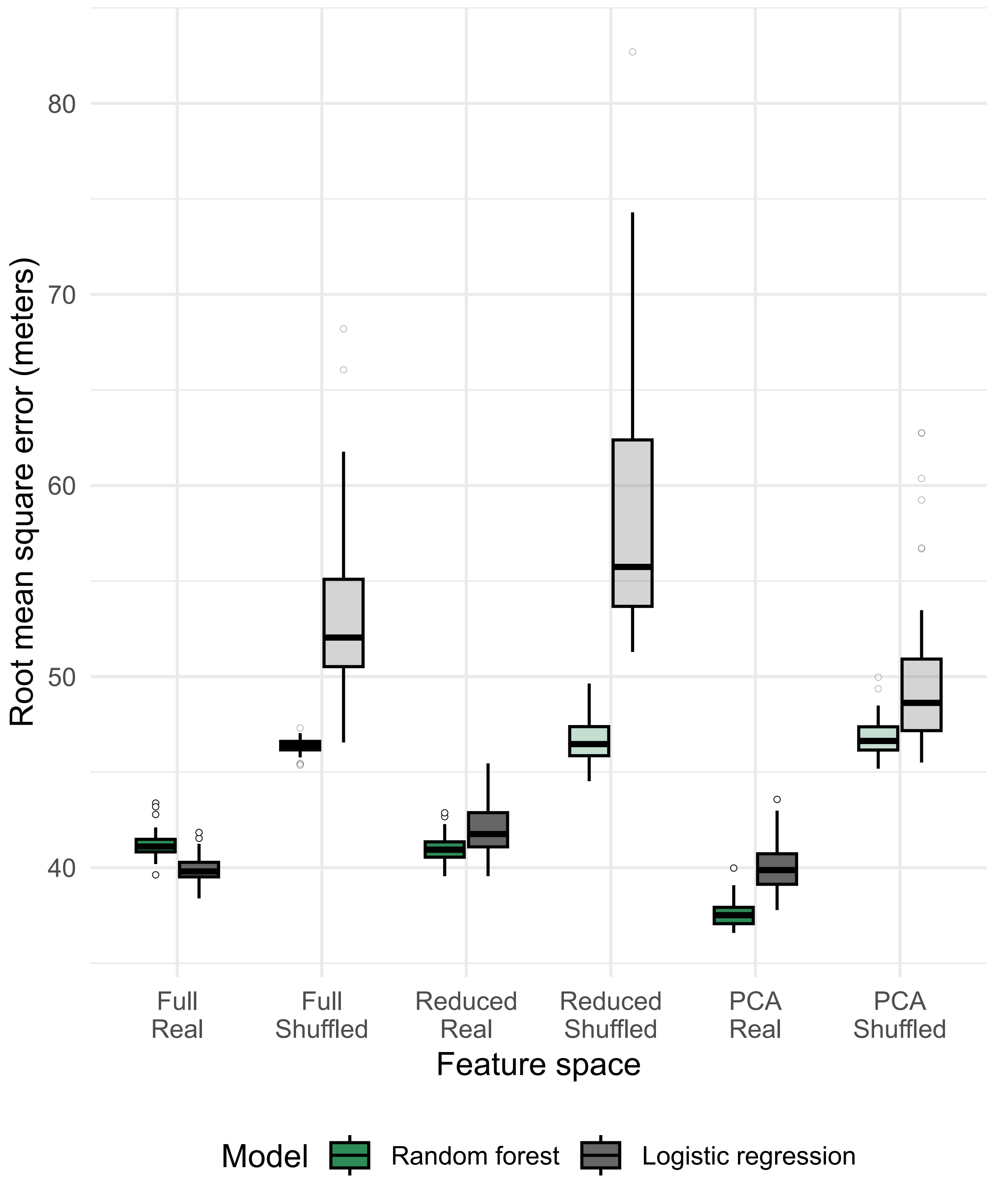


**Figure S2. Depth of isolation performance vs. shuffled label baselines.**

For the two algorithms chosen (logistic regression and random forest) and three feature spaces used (full pangenome, reduced pangenome, PCA components), we compared classification performance of all folds and resamples runs for the models trained on real data (depth of isolation labels taken from JGI and NCBI metadata) and data where the depth of isolation labels were randomly shuffled. For real models (darker colored boxes), the root mean square error (RMSE) ranges from 37.3m–41.3m. For shuffled models (lighter colored boxes), RMSE values were higher, at 46.4m–58.7m. While real models did outperform these randomized baselines, their RMSE values were very high relative to the overall range of depth of isolation values. Any signal detected in these models appears weak.

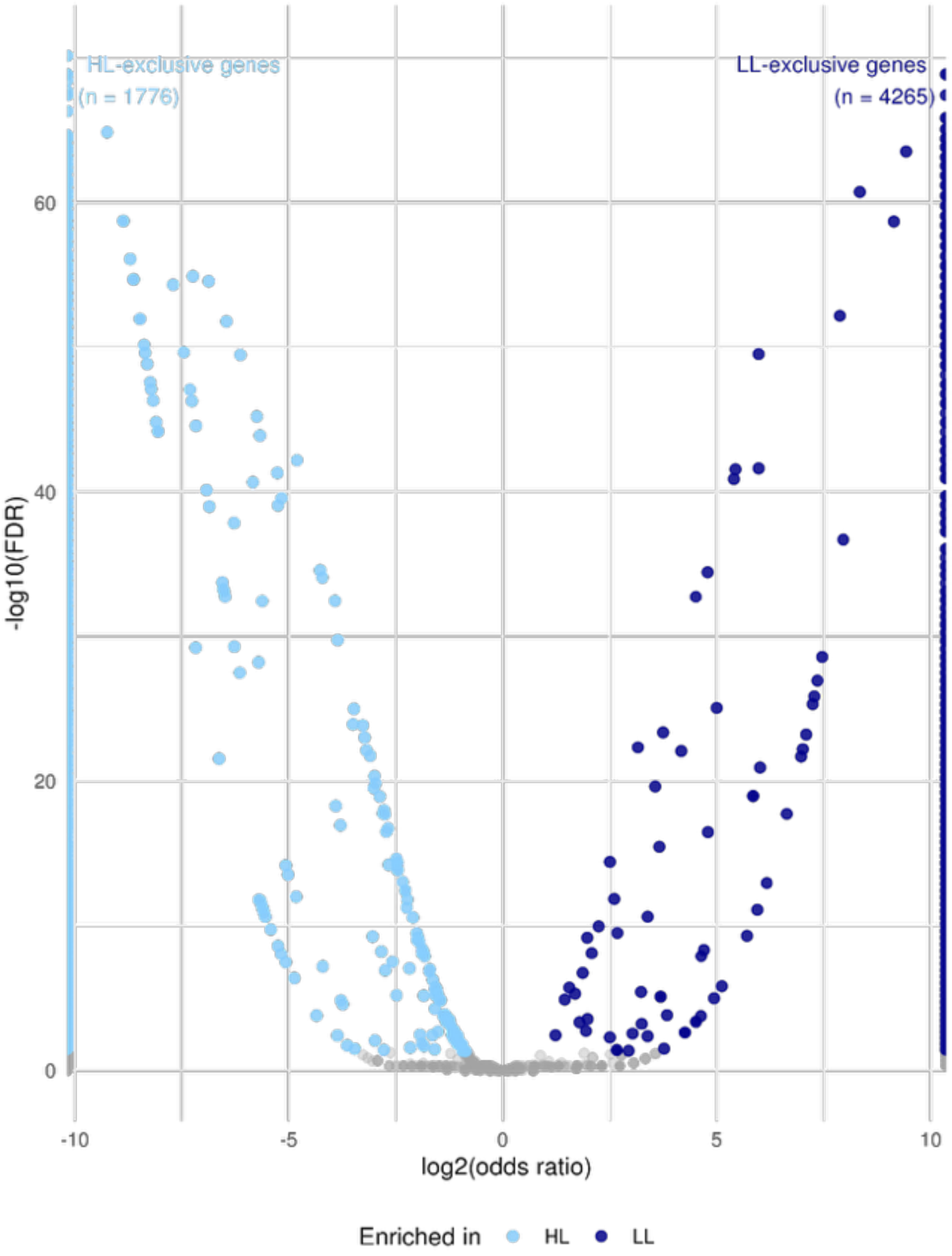


**Figure S3. Volcano plot of differentially encoded genes across ecotypes.**

A substantial fraction of genes was significantly differently distributed between HL/LL ecotypes. Fisher testing with multiple testing correction applied yields thousands of significant genes (**Table S2**), with many of them being mutually exclusive to one ecotype or the other (flanking left and right pillars at OD = +/- 10). For genes shared across ecotypes, 174 were significantly enriched in HL (left), and 68 were significantly enriched in LL (right). These numbers should be interpreted with caution, as sequence-based gene clustering can result in canonical and variant clusters of the same gene. As such, some genes and functional roles may still be conserved across ecotypes, but the orthologs differ to such an extent that they are classified as separate gene clusters (e.g., see pepA text in Results). Furthermore, variable completeness of these genomes (measured by BUSCO score) means many genes may be missed by this high-level analysis due to fundamental incompleteness in public data.

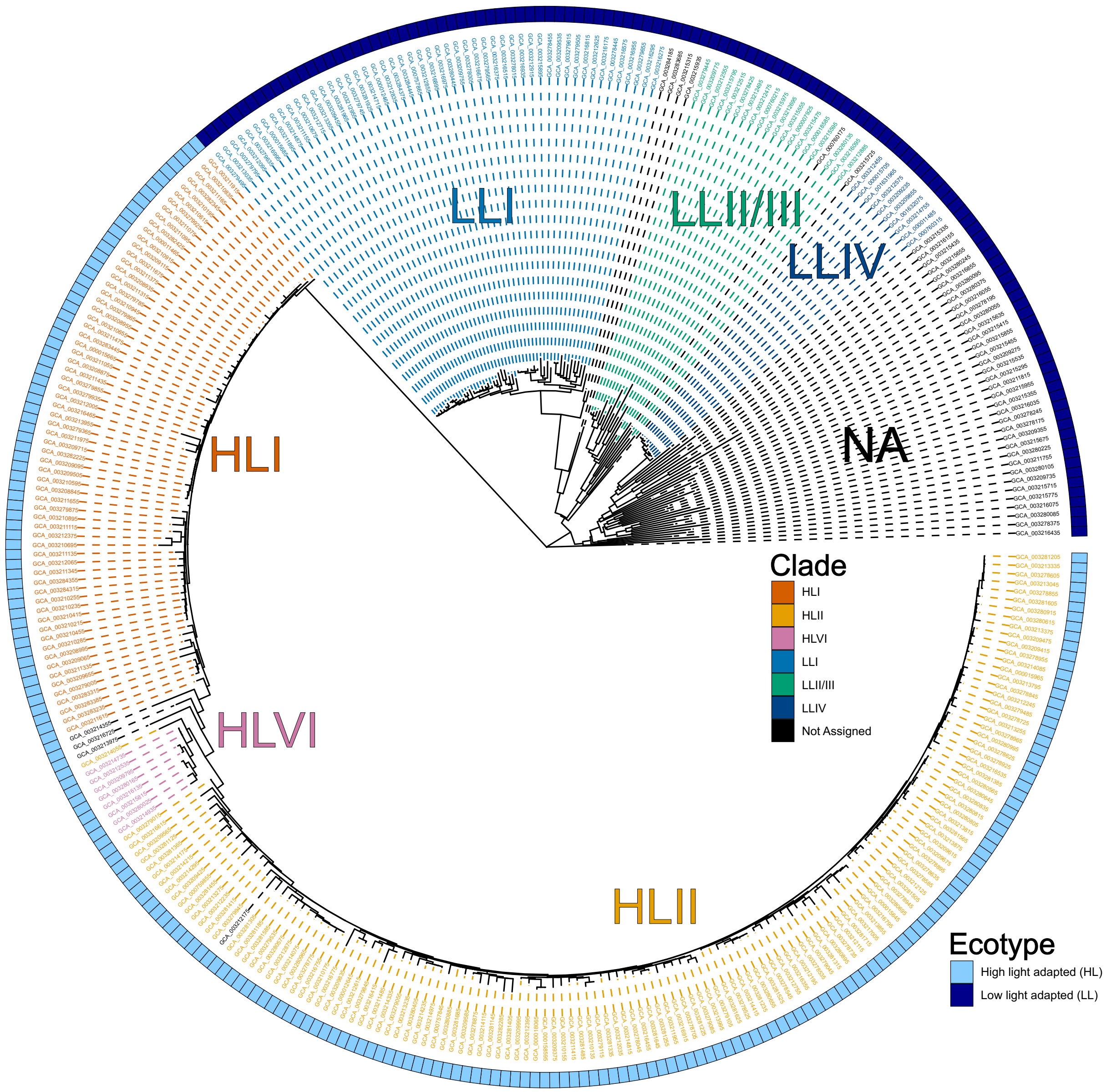


**Figure S4. Internal transcribed spacer-based phylogeny.**

We extracted the sequence between 16s and 23s rRNA genes from genomes with sufficient sequence coverage and ecotype labels, aligned them, and built a maximum likelihood phylogenetic tree to see if clades could be called with adequate statistical support. Our bootstrap support for the HL/LL ecotype split (outer ring, colored boxes) was strong, but nearly all other clades had minimal support (bootstrap support < 50) that prevented us from using the tree for clade-calling. However, preassigned clade labels from ProPortal were present for most genomes with ecotype data, and these are overlaid by coloring the tip labels. These clades map cleanly to the overall tree structure.

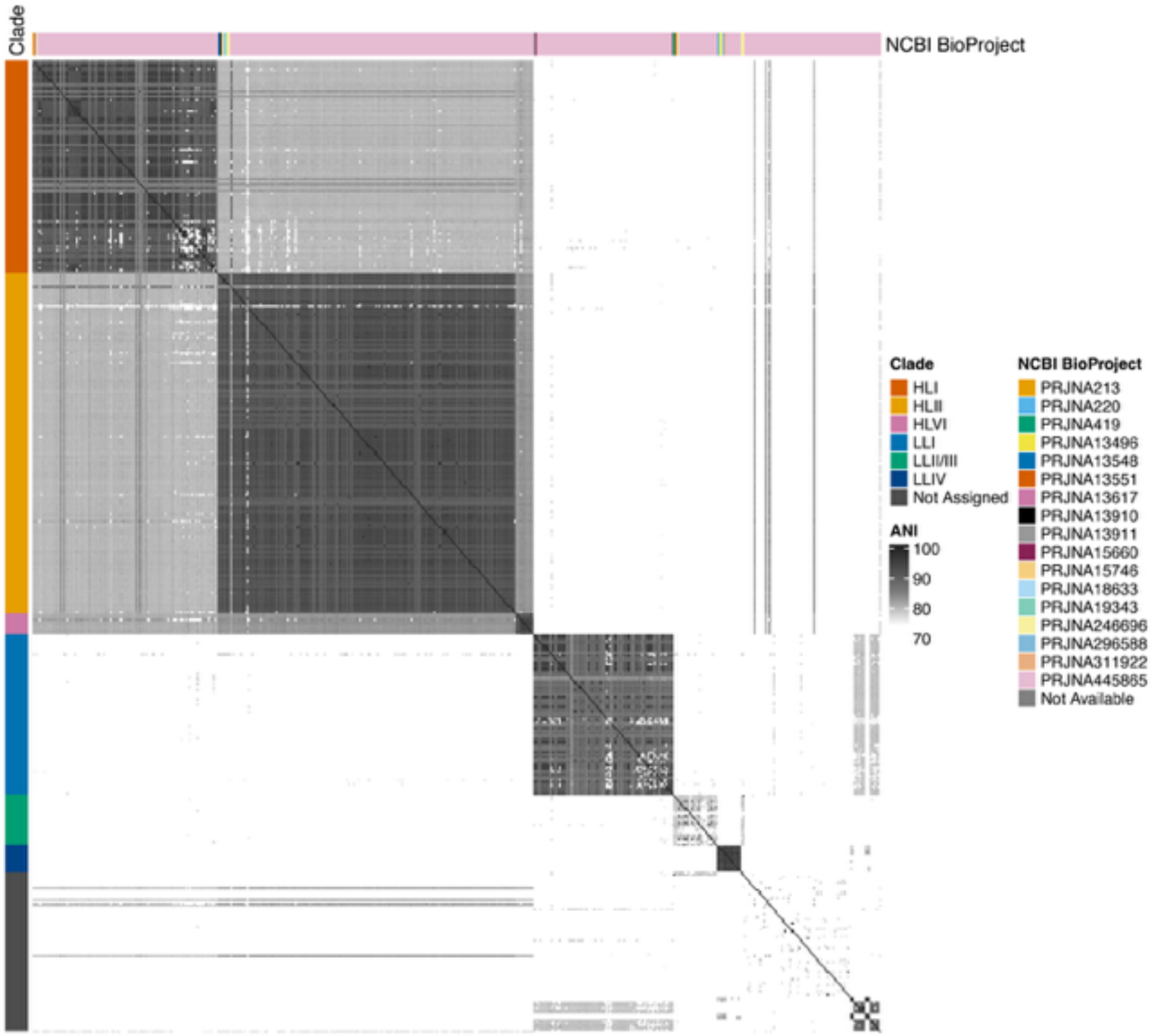


**Figure S5. BioProject-associated batch effects do not drive grouping.**

Given the robust grouping of FastANI-based identity by assigned clades, we plotted ANI ordered by clade but then sorted by the associated NCBI BioProject ID. The clade-based blocks remain predominant, and we do not see signs that batch effects that are routinely introduced in any sequencing project are driving any of the clustering observed in our project. BioProject 445865 (data by Berube et al.) dominate the samples used in our dataset, and were specifically collected in their study with a focus on expanding the number of sequences available from oceans around the world. Other BioProject-associated samples do not show patterns distinct from those seen in this dominant BioProject, supporting that true biological diversity is represented in the blocks.

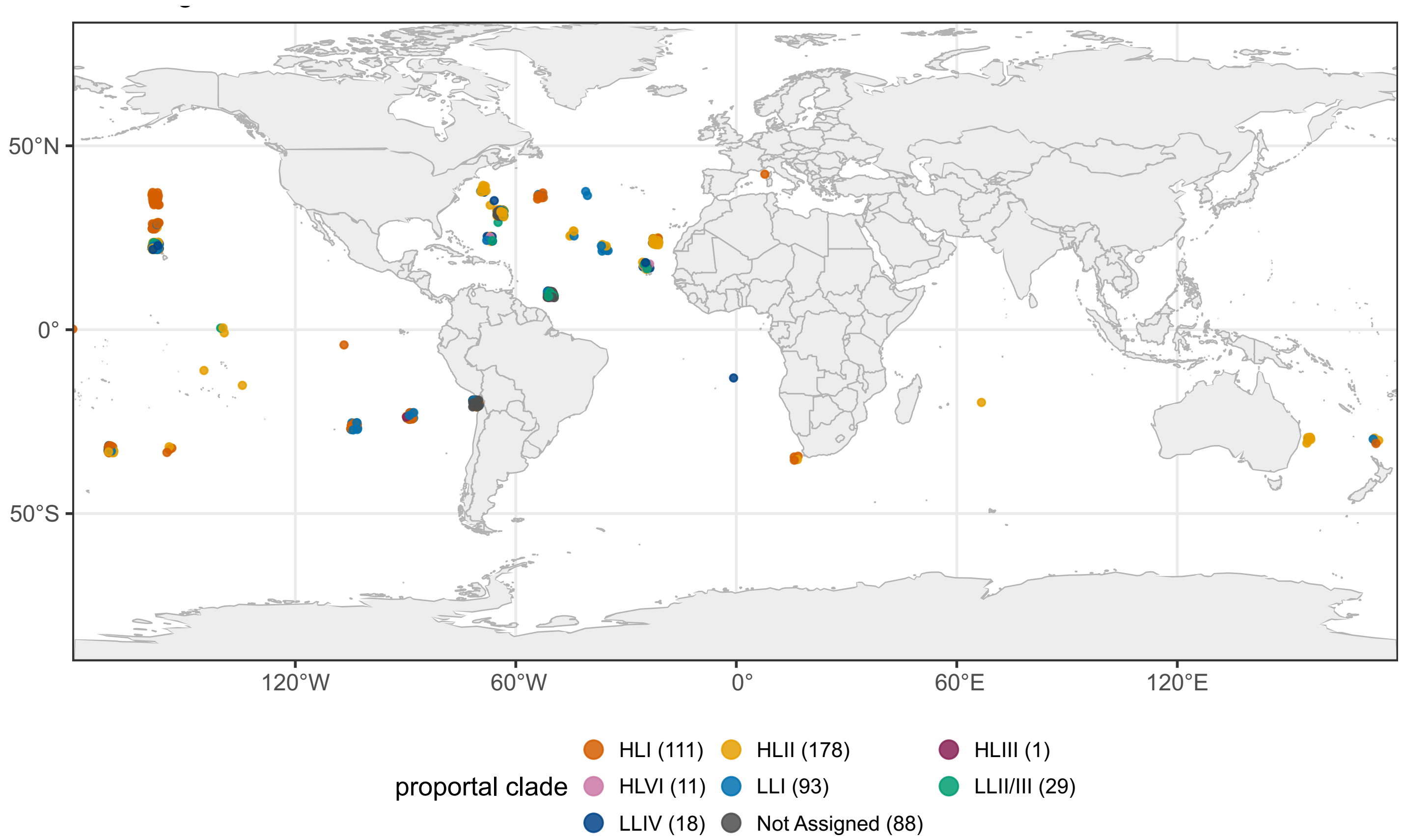


**Figure S6. Global distribution of samples by assigned clade.**

While isolates and clades are found with a global distribution, the vast majority of samples tend to be found in tightly clustered sites, specifically in the northern Atlantic and Pacific oceans, with other clusters in the southern and eastern Pacific. The diversity seen in genomic data is additionally unsurprising given the sheer distances observed even between sample clusters in single ocean basins. Latitude and longitude of points on this map have been randomly jittered by ±1° from their true values to illustrate density of clusters.

### Supplementary Tables

**Table S1. Shuffled model performance correlations.**

To assess whether high variation in per repeat performance in the shuffled label baseline models, we performed Spearman and Pearson correlation for the per repeat MCC values and the proportion of shuffled label isolates that were randomly assigned to the correct label. Across models, all correlation values are high.

**Table S2. Features identified in ML modeling for ecotype and depth.**

The set of identified ML model features across model types. The specific sheets are:

*LR_top*: The top 50 features from full feature space logistic regression (LR) models predicting ecotype.

*LR_features*: The FASTA formatted protein sequences representative of each PanTA gene cluster for these top LR features.

*LR-RF_intersect*: The top features identified both LR and random forest (RF) models.

*Intersect_features*: The FASTA formatted protein sequences representative of each PanTA gene cluster for the top features shared between LR and RF models.

*RF_top*: The top 37 features from full feature space random RF predicting ecotype. The feature space showed far greater variability for RF models, and only 37 features occurred across multiple RF iterations.

*RF_features*: The FASTA formatted protein sequences representative of each PanTA gene cluster for these top RF features.

*Depth_top*: The top features identified by the best performing depth model (PCA-reduced feature space RF) across iterations. We extracted the 50 features that made up principal component 1, the top predictor.

*Depth_features*: The FASTA formatted protein sequences representative of each PanTA gene cluster for these top PCA RF features.

**Table S3. Fisher's test results for PanTA gene cluster distribution across ecotype.**

Fisher's testing with adjustment for multiple comparisons across gene clusters. Many genes show statistically significant distribution differences across ecotypes.

**Table S4. Differences in genome metrics across Prochlorococcus clades.**

After Kruskal-Wallis testing supported significance differences between genome composition metrics across clades, we performed Dunn testing with Benjamini-Hochberg correction to identify specific pairwise comparisons that differed across clades. Small total sample sizes for some clades limited statistical confidence.
